# Over eight hundred cannabis strains characterized by the relationship between their psychoactive effects, perceptual profiles, and chemical compositions

**DOI:** 10.1101/759696

**Authors:** Alethia de la Fuente, Federico Zamberlan, Andrés Sánchez Ferrán, Facundo Carrillo, Enzo Tagliazucchi, Carla Pallavicini

## Abstract

**Background:** Commercially available cannabis strains have multiplied in recent years as a consequence of regional changes in legislation for medicinal and recreational use. Lack of a standardized system to label plants and seeds hinders the consistent identification of particular strains with their elicited psychoactive effects. The objective of this work was to leverage information extracted from large databases to improve the identification and characterization of cannabis strains.

**Methods:** We analyzed a large publicly available dataset where users freely reported their experiences with cannabis strains, including different subjective effects and flavour associations. This analysis was complemented with information on the chemical composition of a subset of the strains. Both supervised and unsupervised machine learning algorithms were applied to classify strains based on self-reported and objective features.

**Results:** Metrics of strain similarity based on self-reported effect and flavour tags allowed machine learning classification into three major clusters corresponding to *Cannabis sativa*, *Cannabis indica*, and hybrids. Synergy between terpene and cannabinoid content was suggested by significative correlations between psychoactive effect and flavour tags. The use of predefined tags was validated by applying semantic analysis tools to unstructured written reviews, also providing breed-specific topics consistent with their purported medicinal and subjective effects. While cannabinoid content was variable even within individual strains, terpene profiles matched the perceptual characterizations made by the users and could be used to predict associations between different psychoactive effects.

**Conclusions:** Our work represents the first data-driven synthesis of self-reported and chemical information in a large number of cannabis strains. Since terpene content is robustly inherited and less influenced by environmental factors, flavour perception could represent a reliable marker to predict the psychoactive effects of cannabis. Our novel methodology contributes to meet the demands for reliable strain classification and characterization in the context of an ever-growing market for medicinal and recreational cannabis.

## Background

*Cannabis indica* and *Cannabis sativa* have been used in traditional medicine for millennia around the world, as well as a source for fiber and food (Mechoulam, 2019; Russo, 2011; Russo et al., 2008). In the past century, Western medicine has gone a long way to find specific medications for many afflictions traditionally treated with cannabis-derived products, and the recreational use of marihuana for its psychoactive properties emerged as the main motivation for its cultivation and consumption (Clarke & Merlin, 2013). This and other factors led to the inclusion of cannabis as a Schedule 1 controlled substance, a category reserved for compounds with “no currently accepted medical use” according to the Food and Drug Administration (U.S. Food and Drug Administration, 2015), despite its long history in the treatment of diverse illnesses, symptoms and conditions (Clarke & Merlin, 2013).

Recently, regional changes in legislation made marihuana and other cannabis-derived products available for medicinal and recreational use (Bifulco & Pisanti, 2015; Fischer, Ala-Leppilampi, Single, & Robins, 2010; Graham, 2015; Hetzer & Walsh, 2014). It is expected that through resilient patient/consumer activism and increasing scientific evidence supporting the medicinal use of cannabis, this phenomenon will continue to rise gradually in more countries (Hazekamp, Tejkalová, & Papadimitriou, 2016). Market growth for marihuana has been dramatic in some countries; for instance, in the United States sales reached $6.7 billion in 2016, with 30% growth year-over-year, representing the second largest cash crop, with total worth over $40 billion (Adams, 2019; Robinson, 2017). These sudden changes created novel problems for users, as cannabis cultivators transition towards legal business models, yet without a world-wide standard for their products. Cannabis dispensaries offer dry cannabis flowers or buds (Gilbert & DiVerdi, 2018), extracts and essential oils (Permanente & Care, 2008) and various edibles (Weedmaps, n.d.); however, since in most countries these products remain illegal, there are no international agreements to regulate their quality or chemical content.

The development of standards is further complicated by the heterogeneous chemical composition inherent to cannabis. Plants contain over 400 compounds, including more than 60 cannabinoids, the main active molecules being tetrahydrocannabinol (THC) and cannabidiol (CBD). These two cannabinoids were often considered the only chemicals involved in the medicinal properties and psychoactive effects associated with cannabis, and remain the only ones screened when evaluating strain chemotypes (De Meijer, Hammond, & Sutton, 2009; Fetterman et al., 1971; Hazekamp et al., 2016; Nie, Henion, & Ryona, 2019; UNODC, 1968). However, increasing evidence supports the relevance of terpenes and terpenoids, molecules responsible for the flavour and scent of the plants, both as synergetic to cannabinoids and as active compounds by themselves (Henry, 2017; Hillig, 2004; Nuutinen, 2018; Russo, 2011). Flavours have predictive value at strain level (Gilbert & DiVerdi, 2018) that may not be superseded by the determination of the species, or even by quantification of THC and CBD content (Jikomes & Zoorob, 2018). Terpenes are widely used as biochemical markers in chemosystematics studies to characterize plant samples due to the fact that they are strongly inherited and relatively unaffected by environmental factors (Aizpurua-Olaizola et al., 2016; Casano, Grassi, Martini, & Michelozzi, 2011; Hillig, 2004). Cannabinoid content, on the other hand, can vary greatly among generations of the same strain, and also due to the sex, age and part of the plant (Fetterman et al., 1971; Hazekamp et al., 2016).

In this work, we combined different sources of data for the classification of cannabis strains, linking both self-reports of psychoactive effects and flavour profiles with information obtained from experimental assays of cannabinoid and terpene content. Our analysis comprised 887 different strains and was based on a large sample (>100.000) of user reviews publicly available at the website Leafly (www.leafly.com). The reports contained unstructured written reviews of experiences for each strain, as well as structured tags indicating flavour profiles and subjective effects. We explored the following four hypotheses: 1) supervised and unsupervised machine learning algorithms can group strains into clusters of similar breeds based on subjective effect tags, but also based on flavour profile tags, 2) certain pairs of effect and flavour tags are correlated across strains, suggesting synergistic effects, 3) unstructured written reports contain information consistent with the tags, and the detection of recurrent topics in the reports matches the known effects and uses of different cannabis breeds, and 4) terpene profiles are consistent with the perceptual characterizations made by the users.

## Methods

### User reported data

Data corresponding to >1.200 cannabis strains was accessed and downloaded from Leafly (www.leafly.com) (August 2018). Leafly is the largest cannabis website in the world wide web, allowing users to rate and review different strains of cannabis and their dispensaries. Sets of predefined tags related to psychoactive effects (e.g. “aroused”, “creative”, “euphoric”, “relaxed”, “paranoid”) and flavours (e.g. “apple”, “coffee”, “flowery”, “apricot”, “vanilla”) are assigned to strains via crowdsourcing, together with a large number (>100.000) of unstructured written reviews. Strains with less than 10 reviews were discarded, resulting in 887 strains included in this study. Each strain was classified as indica, sativa or hybrid. Users associate strains with tags indicating flavours (48 different tags) and effects (19 different tags). Details on all included strains, flavour and effect tags are presented in additional tables [see Additional file 1].

### Effect and flavour tags

Given a strain s with n reviews, we considered for the i-th review the vectors 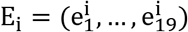 and 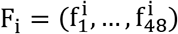 where 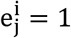 if the tag for the j-th effect appeared in the i-th review, and 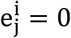 otherwise. The 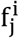 were defined analogously, but based on the flavour tags. Next, the strain was identified with the vectors 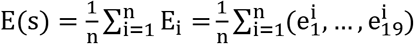 and 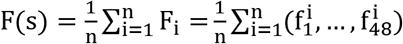, representing the probability that each effect and flavour tag was used in the description of the strain.

### Network and modularity analysis

Given two strains s_1_ and s_2_, they were represented in the effect / flavour network by nodes linked with a connection weighted by the value of the non-parametric Spearman correlation between vectors E(s_1_) and E(s_2_) / F(s_1_) and F(s_2_), respectively. To find sub-networks with dense internal connections and sparse external connections (i.e. modules), the Louvain agglomerative algorithm (Blondel, Guillaume, Lambiotte, & Lefebvre, 2008) was applied to maximize Newman’s modularity using a resolution parameter γ = 1. To visualize the resulting networks, we used the ForceAtlas 2 layout included in Gephi (Bastian, Heymann, & Jacomy, 2009) (https://gephi.org/). ForceAtlas 2 represents the network in two dimensions, modeling the link weights (i.e. Spearman correlations) as springs, and the nodes as point charges of the same sign. The attraction is then computed using Hook’s law (Hooke, 1678) and the repulsion using Coulomb’s law (Coulomb, 1785).

### Effect-flavour correlation analysis

For all strains, the effect and flavour frequency vectors can be summarized as matrices E_is_ and F_is_of size 887×19 and 887×48, respectively, indicating the probability of observing the i-th effect / flavour tag in strain s. To investigate associations between effect and flavour tags, we computed all 19×48 = 912 non-parametric Spearman correlations between all possible pairs of columns from 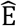 and 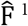. Since some effect and flavour tags appeared very sparsely, we only considered pairs of strains for which at least one report included the given flavour / effect tag (i.e. we excluded columns of zeros from the correlation analysis).

### Random forest classifiers

To investigate whether effect and flavour tags could discriminate between different cannabis species, we trained and evaluated (5-fold stratified cross-validation) machine learning classifiers to distinguish the 265 indicas from the 171 sativas in the dataset, using as features the corresponding E(s) and F(s) vectors for each strain s.

Classifiers were based on the random forest algorithm, as implemented in scikit-learn (https://scikit-learn.org/). This algorithm builds upon the concept of a decision tree classifier, in which the samples are iteratively split into two branches, depending on the values of their features. For each feature, a threshold is determined so that the samples are separated to maximize a metric of the homogeneity of the class labels assigned to the resulting branches. The algorithm stops when a split results in a branch where all the samples belong to the same class, or when all features are already used for a split. This procedure is prone to overfitting, because a noisy or unreliable feature selected early in the division process could bias the remaining part of the decision tree. To attenuate this potential issue, the random forest algorithm creates an ensemble of decision trees based on a randomly chosen subset of the features. After training each tree in the ensemble, the probability of a new sample belonging to each class is determined by the aggregated vote of all decision trees. We trained random forests using 1.000 decision trees and a random subset of features of size equal to the rounded square root of the total number of features. The quality of each split in the decision trees was measured using Gini impurity, and the individual trees were expanded until all leaves were pure (i.e. no maximum depth was introduced). No minimum impurity decrease was enforced at each split, and no minimum number of samples were required at the leaf nodes of the decision trees. All model hyperparameters are detailed in the scikit-learn documentation (https://scikit-learn.org/).

To assess the statistical significance of the output, we trained and evaluated 1.000 independent random forest classifiers using the same features but after scrambling the class labels. We then constructed an empirical p-value by counting the number of times that the accuracy of the classifier based on the scrambled labels exceeded that of the original classifier. The accuracy of each individual classifier was determined by the area under the receiver operating characteristic curve (AUC).

### Natural language processing of written unstructured reports

Text preprocessing was performed using the Natural Language Toolkit (NLTK, http://www.nltk.org/) in Python 3.4.6. The following steps were applied: 1) discarding all punctuation marks (word repetitions allowed) and splitting into individual words, 2) word conversion to the root from which the word is inflicted using NLTK (i.e. lemmatization), 3) conversion to lowercase, 4) after lemmatization, words containing less than two characters were discarded (Sanz & Tagliazucchi, 2018).

To quantitatively explore the semantic content of the reports we used Latent Semantic Analysis (LSA) (Landauer, Foltz, & Laham, 1998; Martial et al., 2019; Sanz & Tagliazucchi, 2018) over all combined strain reports (N = 100.901) in the subsamples of indicas (30.977 reports from 265 strains) and sativas (18.193 reports from 171 strains). For this, we constructed a matrix A_ij_ containing in its i,j position the weighted frequency of the i-th term in the combined reports of the j-th strain. The weighted frequency (tf-idf weighting) was computed as 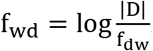, where f_wd_represents the frequency of word w in document d, |D| indicates the total number of documents, and f_dw_ is the fraction of documents in which word w appears. To avoid very common / uncommon words, we kept only those appearing in more than 5% / less than 95% of the documents, respectively.

LSA was applied to reduce the rank of A_ij_, thus reducing its sparsity and identifying different words by semantic context. For this purpose, the matrix was first decomposed using Singular Value Decomposition (SVD) into the product of three matrices (Huang & Narendra, 2008) as 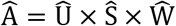, where 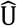 contains the matrix eigenvectors, 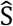 is a diagonal matrix containing the ordered eigenvalues of 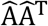, and 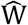 contains the eigenvectors of 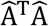 To reduce the dimensionality of the semantic space, only the first 50 singular values of 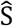 were retained, yielding the truncated diagonal matrix 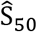. From this matrix, the rank reduced matrix was computed as 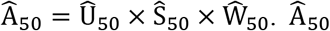 is here referred to as the reduced rank word-document matrix. By computing the Spearman correlation coefficient between the columns of 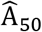 it is possible to estimate the semantic similarity between the written reports associated with pairs of strains. Alternatively, this can be conceptualized as a network, where nodes correspond to strains and links are weighted by the semantic similarity between their associated sets of reports. The choice of rank 50 was validated by investigating the stability of the number of communities and the modularity values detected in this network using the Louvain algorithm. This validation is included as an additional figure [see Additional file 1].

### Principal component analysis and topic detection

To reduce the term-document matrix into a smaller number of components capturing topics appearing recurrently in the corpus of reports, we performed a principal component analysis (PCA) using MATLAB SVD decomposition algorithm. We analyzed the first five components, i.e. the components explaining most of the variance. Each component consisted of a combination of words present in the vocabulary, and the coefficients were used to represent the importance of the words.

### Association between tags and unstructured written reports

To provide an example of the relationship between the reported effect tags and the unstructured written reports, we performed the LSA analysis on two strains with a large number of reports: a strain representative of sativa (Super lemon haze, 1.373 reports), and another representative of indica (Blueberry, 1.456 reports). In this case, the matrix A_ij_ was constructed so that rows represented unique terms in the vocabulary, and columns represented individual reports (i.e. the reports were not pooled for each strain). We then performed PCA for each of the strains and retained the first 25 terms included in the first five components, comparing them afterwards to the most frequently reported effect tags for each strain. The semantic comparison was performed using the Datamuse API (www.datamuse.com), a word-finding engine based on word2vec (Minarro-Gimenez, Marin-Alonso, & Samwald, 2014), an embedding method using shallow neural networks to map words into a vector space with the constraint that words appearing in similar contexts are also close in the vector space embedding. We applied this tool to measure the mean distance of each tag to the words in each component, and then compared this distance to the one obtained using random English words extracted from www.wordcounter.net/random-word-generator.

### Terpene and cannabinoid data

Cannabinoid and terpene profiles of commercial samples of cannabis strains were downloaded from the PSI Labs webpage (www.psilabs.org) (August 2018). This website contains a large number (>1.600) of test results, with mass spectrometry profiles for 14 cannabinoids and 33 terpenes. We downloaded test results corresponding to strains with more than 10 reports in Leafly, yielding a sample of 443 test results from 183 different strains. We discarded terpenes and cannabinoids that were reported in less that three strains, resulting in profiles comprising 10 cannabinoids and 26 terpenes.

## Results

### Effect similarity network

We first investigated the effect similarity network, where each node represented a cannabis strain and links were weighted by the correlation between their effect tag frequency vectors, as defined in the “Effect and flavour tags” section of the Methods. This network is represented in Fig. 1A using the ForceAtlas 2 layout, which increases the proximity of nodes with strong connections. The left panel of Fig. 1A is coloured according to the modules detected using the Louvain algorithm, while the right panel is color-coded based on species (indica, sativa and hybrids). The algorithm produced a partition with modularity Q = 0.221 and a total of 18 modules, of which the largest five contained ≈98% of all strains. The network color-coded by species showed a clear separation of indicas and sativas, with strains labeled as hybrids located in between. Module 1 contained most of the sativa strains, while indicas and hybrids appeared distributed across the other modules.

**Fig. 1.**
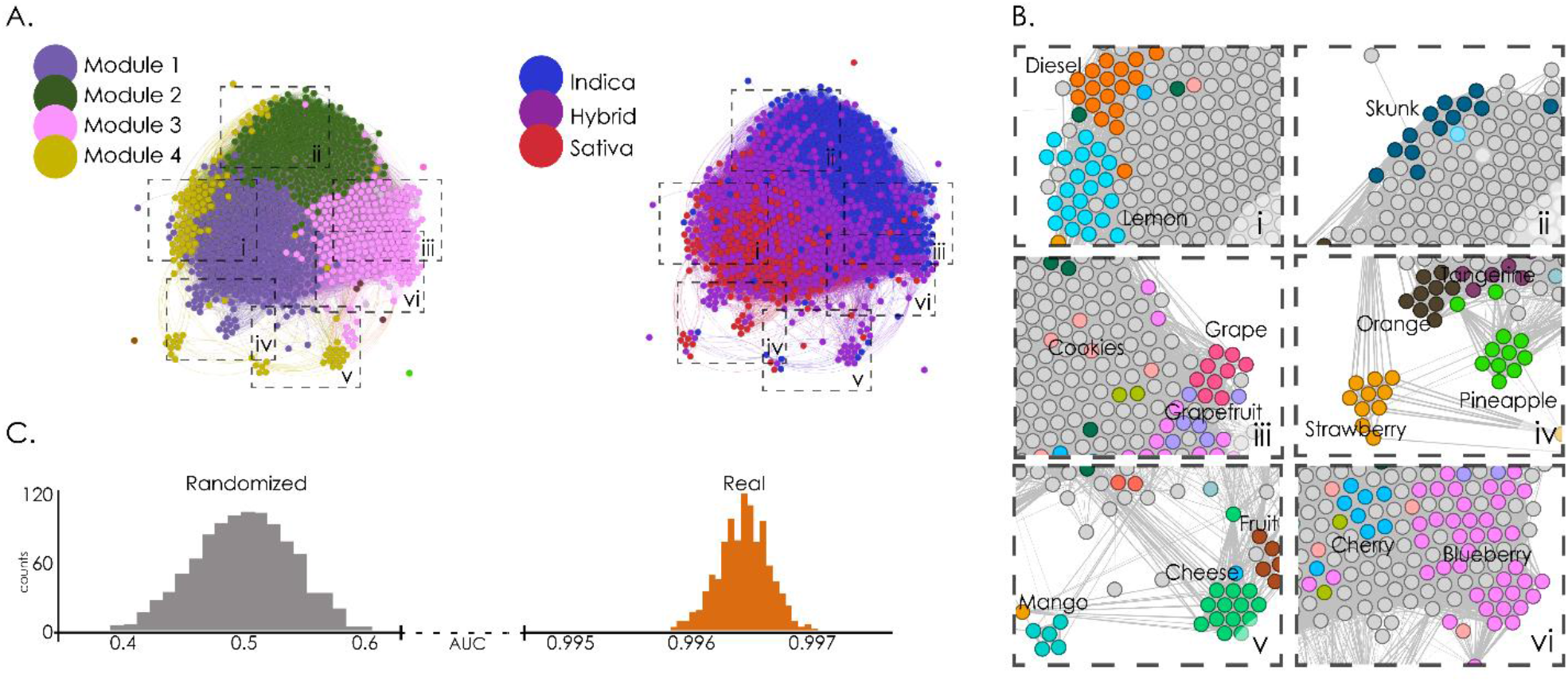
Analysis of the effect similarity network allowed supervised and unsupervised cannabis species classification. A. Effect similarity networks, with nodes representing strains and spatial proximity reflecting the Spearman correlation of the corresponding effect frequency vectors. The left panel is color-coded based on the results of modularity optimization using the Louvain algorithm, while the right panel is color-coded based on species (indica, sativa, hybrids). B. Subpanels zooming into different regions of the networks to show that strains sharing naming conventions were grouped together. C. Histogram of AUC values obtained over 1000 iterations of random forest classifiers using the effect frequency vectors as features and species (indica and sativa) as labels. “Randomized” indicates counts of AUC values obtained after randomly shuffling the sample labels.

Sub-panels I-VI (Fig. 1B) zoom into different regions of the network, showing that strains with similar naming conventions were strongly connected in the effect similarity graph. This was the case for lemons and diesels (I), skunks (II), grapes, cherries and berries (III), pineapples, oranges and strawberries (IV), fruits, cheeses and mangos (V), and blueberries (VI) ^2^. This grouping suggests the presence of correlations between effect and flavour tags, a possibility which is explored in the following sections.

Using the effect tag frequency vectors E(s) as features in a random forest classifier trained to distinguish indicas from sativas resulted in a highly accurate classification (Fig. 1C), with <AUC> = 0.9965 ± 0.0002 (mean ± STD, p < 0.001).

### Flavour similarity network

Next, we studied the network constructed using flavour similarity to weight the links between strains, e.g. the correlation between the F(s) vectors. The resulting network is shown in Fig. 2A (left panel color-coded by modules and right panel color-coded by species). Application of the Louvain algorithm yielded Q = 0.264 and a total of 19 modules, with the four largest containing ≈98% of all strains. In this case, modules comprised predominantly of a single species were no longer clearly visible; however, a gradient of species (from indicas to hybrids to sativas) could be observed from top to bottom.

The amplified panels show that not only strains with similar naming conventions were grouped together, but also that their grouping was related to the flavours represented in their names (Fig. 2B). For instance, blueberries were grouped together and close to a cluster of grapes (I), fruit and cheese strains were in the same subpanel (II), fruit-related strains (pineapple, tangerine, citrus, orange) were grouped together (III), as well as skunks and diesels (IV), mangos and strawberries (V), with lemons appearing cohesively clustered together (VI).

**Fig. 2.**
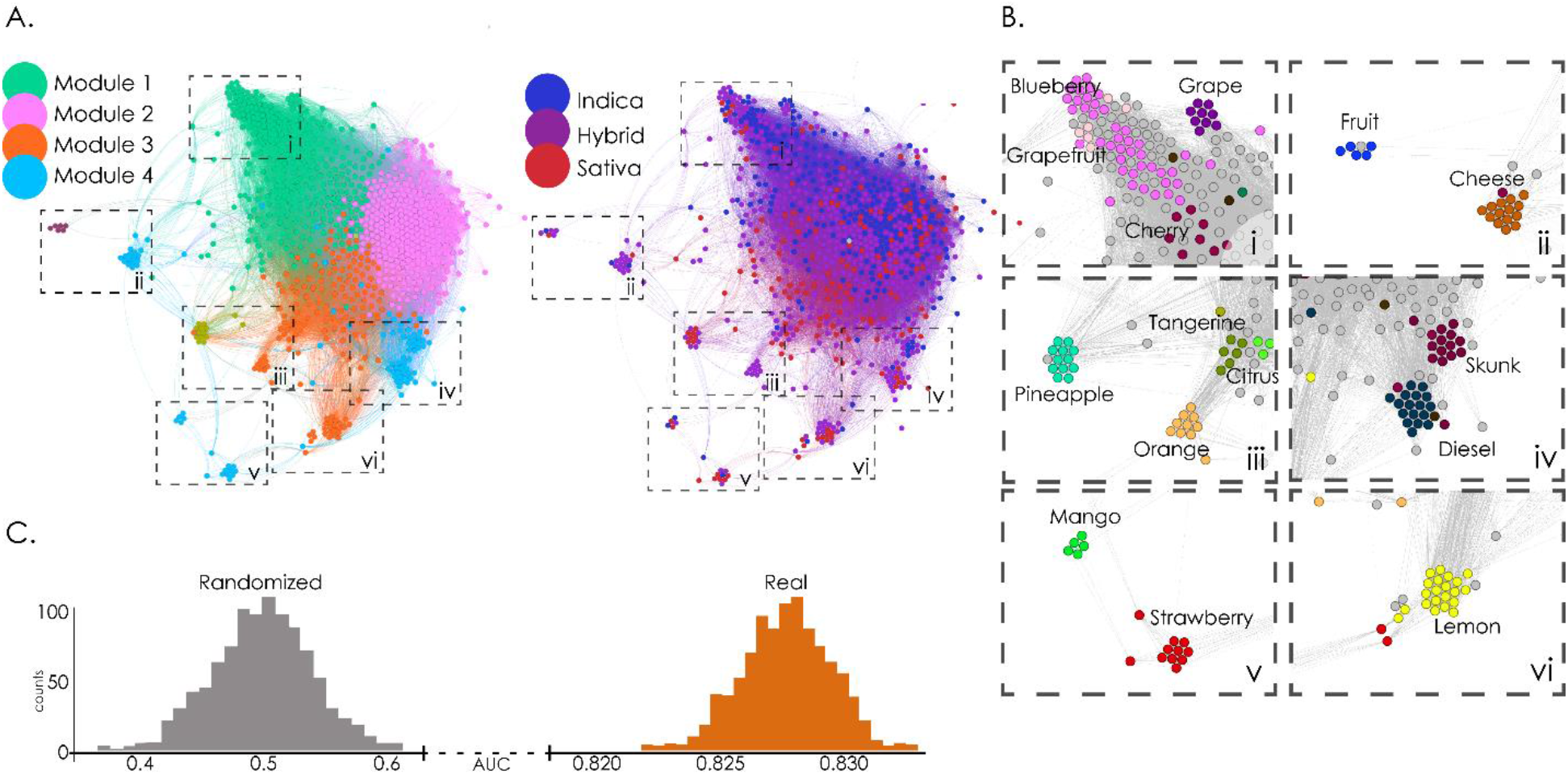
Analysis of the flavour similarity network allowed supervised and unsupervised cannabis species classification. A. Flavour similarity networks, with nodes representing strains and spatial proximity reflecting the Spearman correlation of the corresponding effect frequency vectors. The left panel is color-coded based on the results of modularity optimization using the Louvain algorithm, while the right panel is color-coded based on species (indica, sativa, hybrid). B. Subpanels zooming into different regions of the networks show that strains sharing naming conventions and flavours were grouped together. C. Histogram of AUC values obtained from 1000 iterations of random forest classifiers using the flavour frequency vectors as features and species (indica and sativa) as labels. “Randomized” indicates counts of AUC values obtained after randomly shuffling the sample labels.

Interestingly, when using the flavour tag frequencies as features in a random forest classifier trained to distinguish indicas from sativas, we also obtained a highly accurate classification (Fig. 2C), with <AUC> = 0.828 ± 0.002 (mean ± STD, p < 0.001).

### Correlation between effect and flavour tags

Given that terpenes can be synergetic to cannabinoids (Nuutinen, 2018; Russo, 2011), and noting the distribution of strains presented in Fig. 1, i.e. strains of similar flavour profile appeared close to each other in the network of effect similarity, we computed the correlation between effect and flavour frequencies across all strains, as described in the “Effect-flavour correlation analysis” section of the Methods.

The results are shown in Fig. 3. We found significative (p<0.05, FDR-corrected) negative and positive effect-flavour correlations, represented by the coloured cells in the figure. Fig. 3A shows negative correlations, i.e. inverse relationships between the frequency of the reported effect and flavour tags, while Fig. 3B illustrates positive correlations. Unpleasant effects, such as “anxious”, “dizzy”, “headache” and “paranoid”, correlated negatively with almost all flavours. This could be explained by considering that negative subjective experiences may outweigh flavour or scent perception. Further inspection of Figs. 3A and 3B reveals that certain flavours, such as “berry”, “blueberry”, “earthy”, “pungent” and “woody”, were negatively correlated with stimulant effects, such as “creative” and “energetic”, and at the same time presented positive correlations with soothing effects such as “relaxed” and “sleepy”. Other flavours, such as “citrus”, “lime”, “tar”, “nutty”, “pineapple” and “tropical” presented the opposite behaviour, i.e. they correlated negatively with soothing effects (“relaxed”, “sleepy”) and positively with stimulant effects (“creative”, “energetic”). The fact that the aforementioned flavours presented inverse correlation patterns with respect to opposite psychoactive effects adds support to the validity of this analysis.

**Fig. 3.**
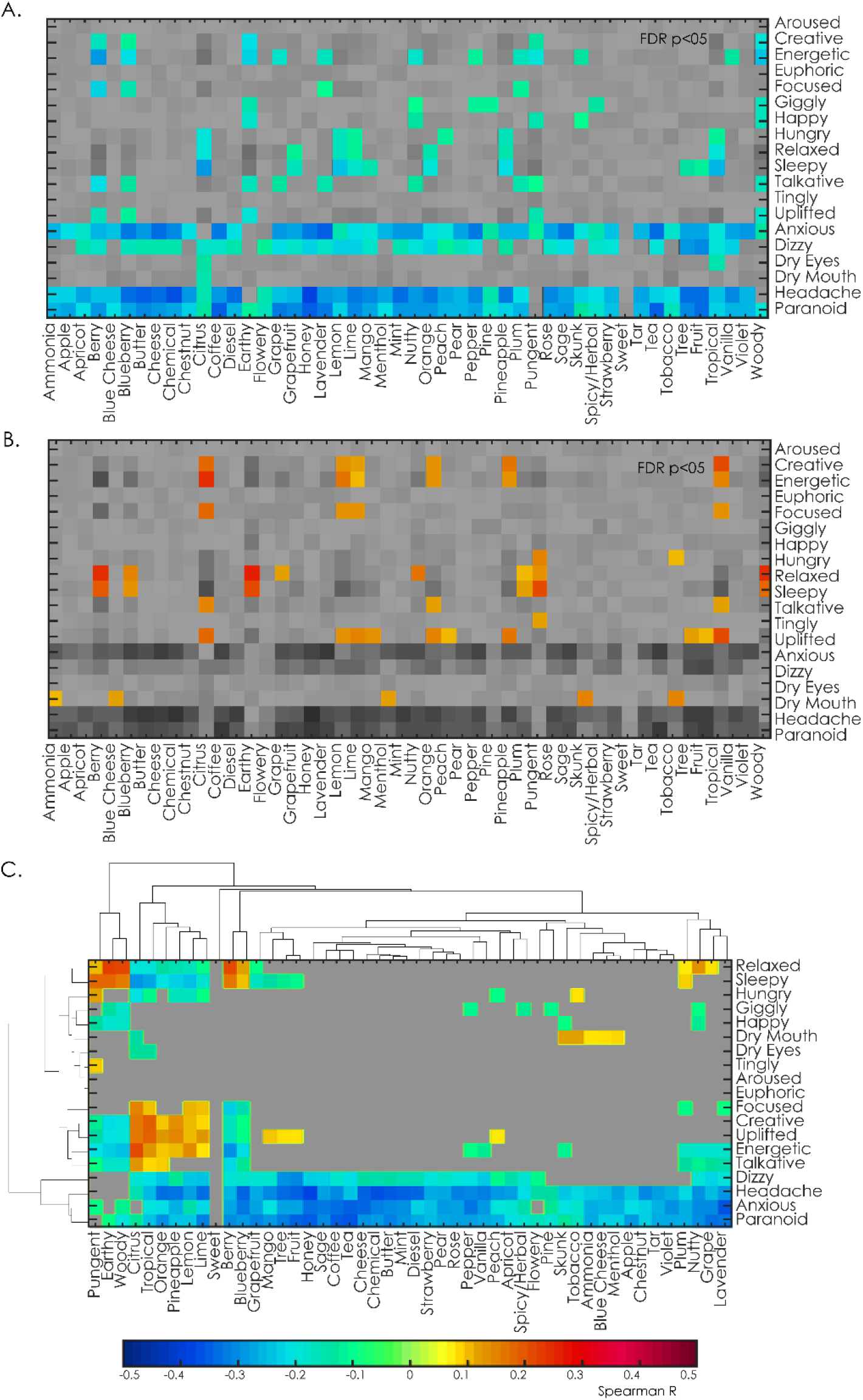
Associations between effects and flavours. A. Significative negative Spearman correlations between effects and flavours. B. Significative positive Spearman correlations between effects and flavours. In both panels significative correlations are indicated using a color scale. Both panels are thresholded at p<0.05, FDR-corrected). C. Hierarchical clustering of the effects and flavours according to their correlations.

Next, we performed a hierarchical clustering of the effects and flavours according to their correlations (Fig. 3C). The main cluster separated unwanted effects from the rest. The remaining clusters of subjective effects were divided into three categories: soothing (“relaxed”, “sleepy”), stimulant (“euphoric”, “creative”, “energetic”, “talkative”) and other miscellaneous effects commonly associated with cannabis use (“hungry”, “giggly”, “happy”, “dry mouth”, “dry eyes”, “tingly” and “aroused”). It is important to note that this hierarchy emerged from considering effect-flavour correlations only. Consistently, flavours were clustered according to their negative correlations (“pungent”, “earthy”, “woody”, “berry”, “blueberry”) and their positive correlations (“citrus”, “tropical”, “orange”, “pineapple”, “lemon”, “lime”).

### Analysis of unstructured written reports

Unstructured written reports can provide complementary information, since users are not limited by predefined sets of effect and flavour tags. We constructed a network in which nodes represented strains and links were weighted by their semantic similarity, measured by the correlation between the columns of the rank-reduced term-document matrix 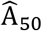 (see the “Natural language processing of written unstructured reports” section in the Methods). The resulting networks are shown in Fig. 4A, where the left panel is color-coded based on module detection and the right panel by species. Applying the Louvain algorithm yielded Q = 0.058, with a total of 15 modules, the largest 4 containing ≈98% of all strains. Module distribution was bimodal, i.e. when compared in terms of unstructured written reports, most strains fell into one of two categories. When comparing the modular decomposition with the species distribution, we found a clear division in terms of indicas and sativas, with hybrids in the border amongst both. This division paralleled the two main modules. Module 1 was conformed predominantly by sativas and hybrids, while module 2 was conformed by indicas and hybrids.

**Fig. 4.**
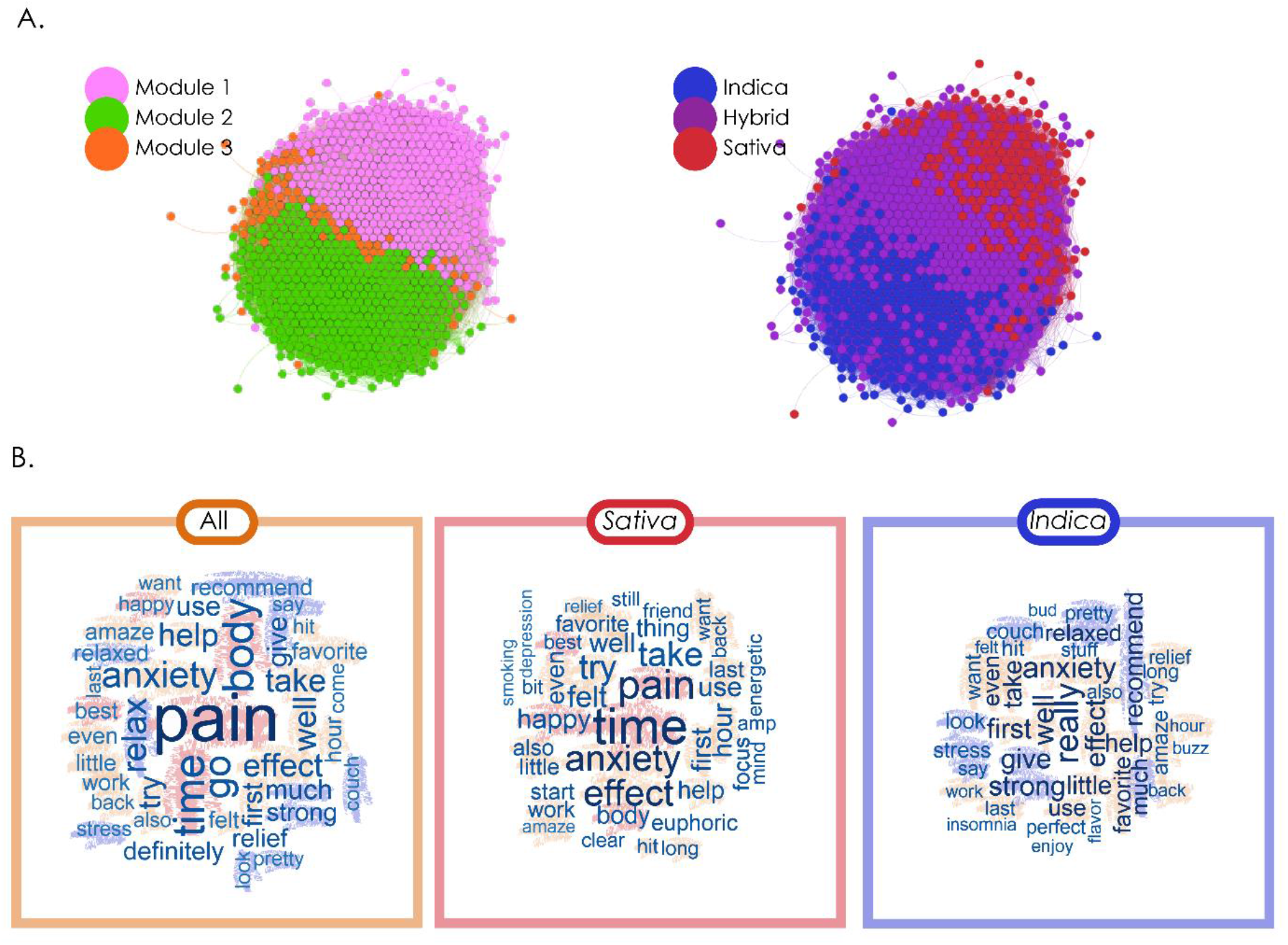
Analysis based on the semantic content of unstructured written reports. A. Networks constructed based on the semantic similarity of the reports associated with strains. The left panel is color-coded based on the results of modularity maximization using the Louvain algorithm, while the right panel is color-coded based on species. B. Word clouds representing the most frequent terms appearing in the reports of all strains combined (left), sativas (middle) and indicas (right).

Next, we investigated the most frequently used terms in the reports of all the strains taken together, and of indicas and sativas considered separately. Fig. 4B presents word cloud representations of the 40 most common terms for all strains combined (left panel), for sativas only (middle panel) and for indicas only (right panel). The most common terms related to the subjective perceptual and bodily effects (terms like “amaze”, “strong”, “felt”, “favourite”, “body”), therapeutic effects and/or medical conditions (“pain”, “anxiety”, “relax”, “help”, “relief”, “focus”) and emotions (“euphoric”, “anxiety”, “happy”, “confusion”). It is important to note that, due to limitations in the amount of available data, this analysis used single term representations (1-grams), therefore words used in positive or negative contexts could not be differentiated, e.g. the term “anxiety” could appear in “This helped calm my anxiety” or in “This caused me anxiety” without distinction.

Half of the most representative words were common to both indicas and sativas, such as “anxiety,” “amaze”, “effect”. The main difference between species emerged after excluding terms common to both, resulting in words such as “focus “, “euphoric“, “energetic” for sativas, and “insomnia“, “enjoy“, “flavour” for indicas. PCA was applied to obtain the main topics present in the written reports. These topics are shown as word clouds in Fig. 5. Upon visual inspection, we found two principal categories of topics: subjective/therapeutic effects, and plant growth/acquisition. The first component obtained from sativa reports consisted of a general mixture of effects, while the second was specific to therapeutic use. For indicas, both components explaining the highest variance related to therapeutic effects. In both cases, the rest of the components were associated to plant growth/acquisition.

**Fig. 5.**
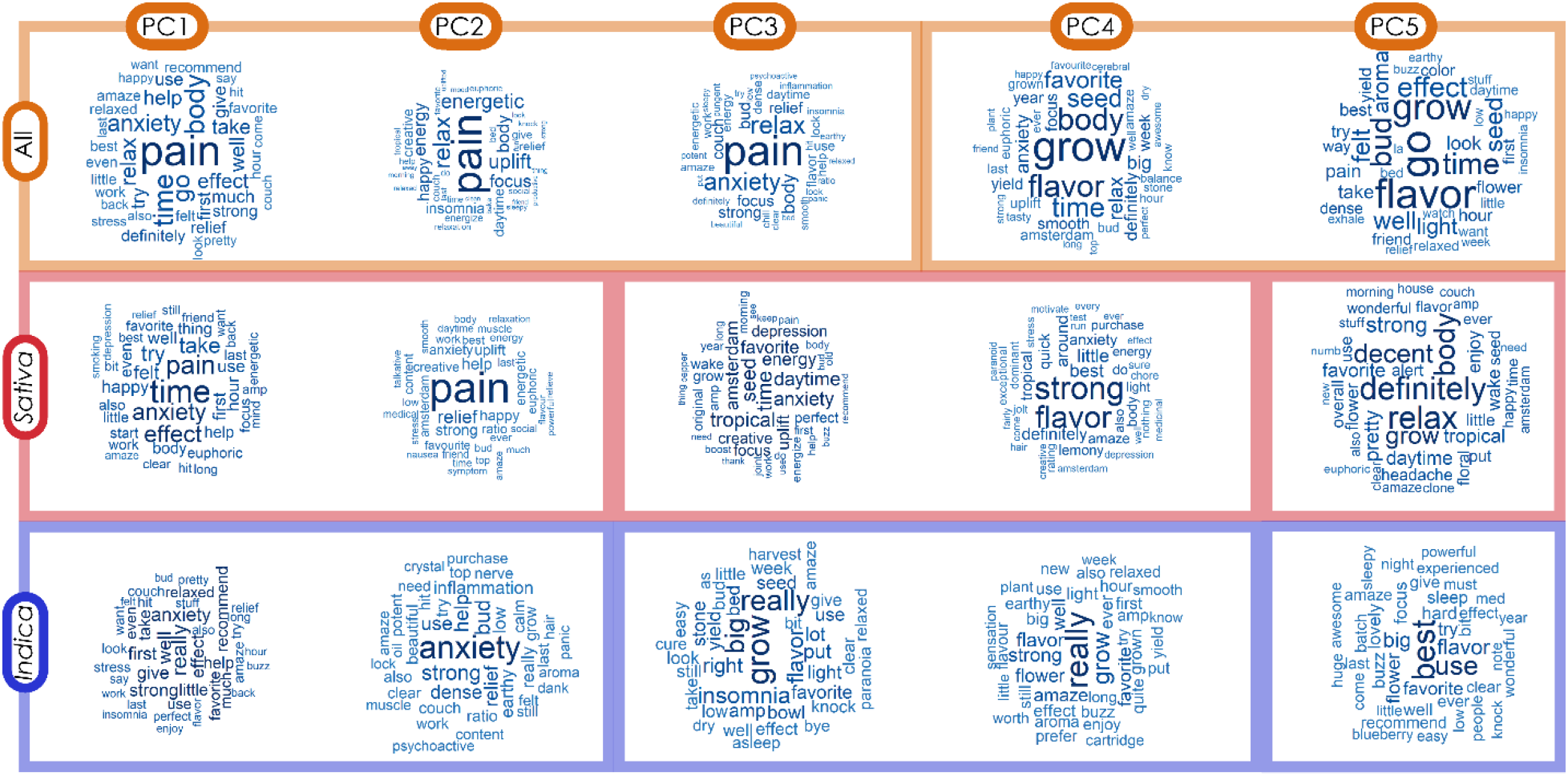
Word clouds representing topics extracted with PCA from the term-document matrix. The first, second and third rows present topics for all strains combined, sativas, and indicas, respectively. Vertical lines separate the content of the topics between subjective/therapeutic effects, and plant growth/acquisition.

To relate the free narrative reports to the effect tags, we investigated two strains with a large number of reports: Super Lemon Haze (sativa, N = 1.373, most frequently reported tags: “happy”, “energetic”, “uplifted”) and Blueberry (indica, N = 1456, most frequently reported tags: “relaxed”, “happy”, “euphoric”). We first applied PCA to the corresponding rank-reduced term-document frequency matrices to obtain the main topics for each strain. The word clouds with the highest-ranking terms for the first 5 principal components of each strain are presented in Fig. 6A. Next, we computed the semantic distance between the most frequent effect tags of each strain and the top 40 words in each of the principal components. The objective of this analysis was to evaluate whether the unstructured written reports reflected the choice of predefined tags made by the users. As shown in Fig. 6B, the most frequently reported effect tags for each strain showed a prominent projection into all the components, as compared to randomly chosen words. This suggests that users selected predefined tags consistently with the contents of their written reports.

**Fig. 6.**
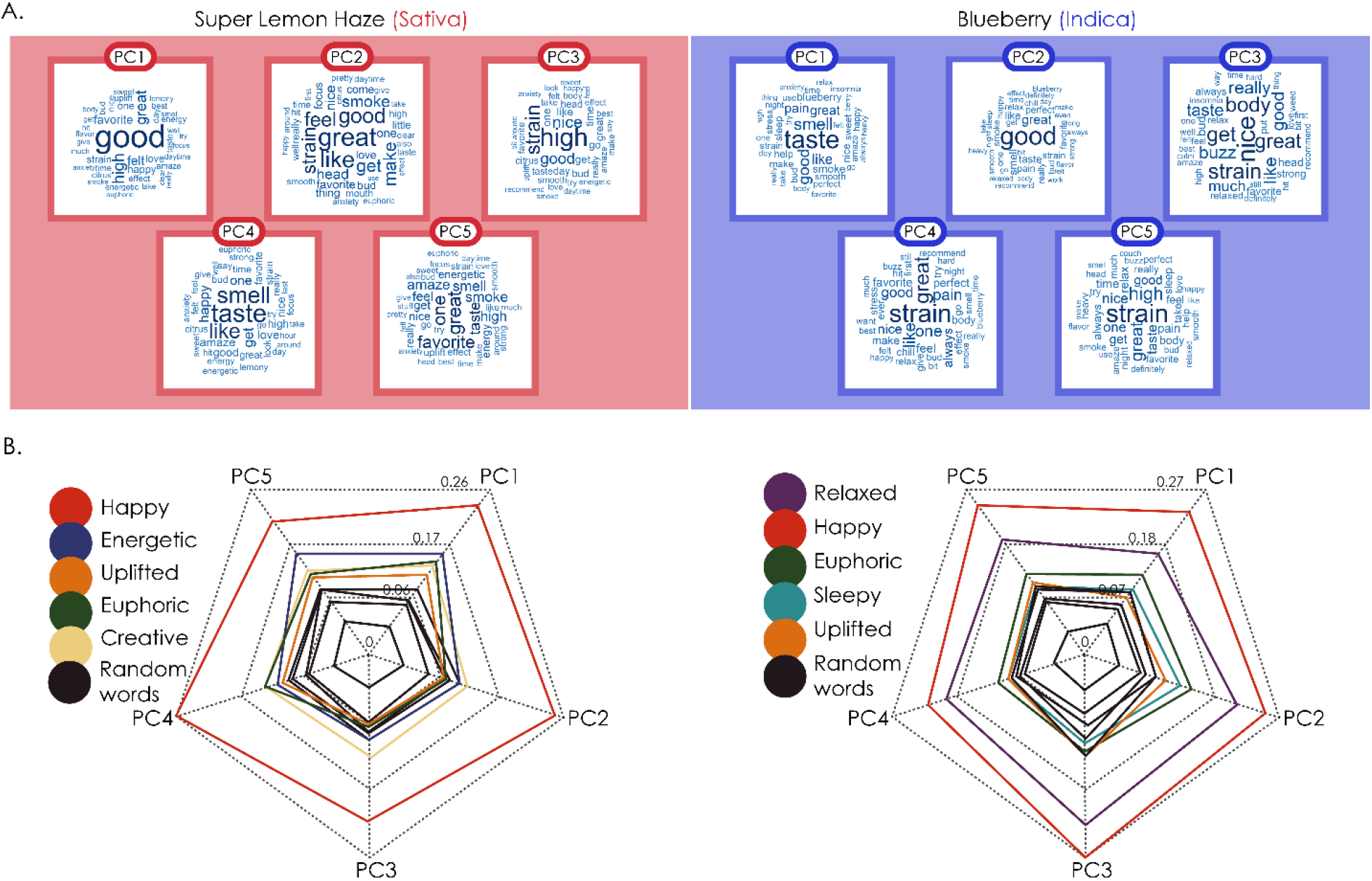
Correspondence between topics extracted from unstructured written reports and the choice of predefined tags. A. Word clouds of the first five principal components for the strains Super Lemon Haze and Blueberry, indicating the most representative topics within the associated reports. B. Radar plots showing the mean semantic similarity between the words in each topic and the most frequently chosen effect tags for both strains. It can be seen that the semantic similarity is the highest for the most frequently used tags, and the lowest for a set of randomly chosen words unrelated to the effects of cannabis.

### Terpene and cannabinoid content

We investigated the relationship between the user reports and the molecular composition of the strains. For this purpose, we accessed publicly available data of cannabinoid content provided in the work of Jikomes and Zoorob (Jikomes & Zoorob, 2018), as well as assays of cannabinoid and terpene content from the PSI Labs website.

The first dataset contains information on THC and CBD content for all 887 strains studied in this work. The relationship between the content of both active cannabinoids is plotted in Fig. 7A, left panel. As reported by Jikomes and Zoorob, the strains fell into three general chemotypes based on their THC:CBD ratios (Jikomes & Zoorob, 2018), consistent with previous findings (Hazekamp et al., 2016; Hillig & Mahlberg, 2004; Jikomes & Zoorob, 2018). Most of the investigated strains fell into chemotype I (Chemotype I: 94.6%, Chemotype II: 4.8%, Chemotype III: 0.6%), indicating high THC vs. CBD ratios. This was replicated using the cannabinoid content data obtained from PSI Labs (N=433 individual flower samples corresponding to 183 different strains), as shown in Fig. 7A, right panel. Again, the majority of the assays corresponded to chemotype I (Chemotype I: 90.3%, Chemotype II: 6%, Chemotype III: 3.7%).

**Fig. 7.**
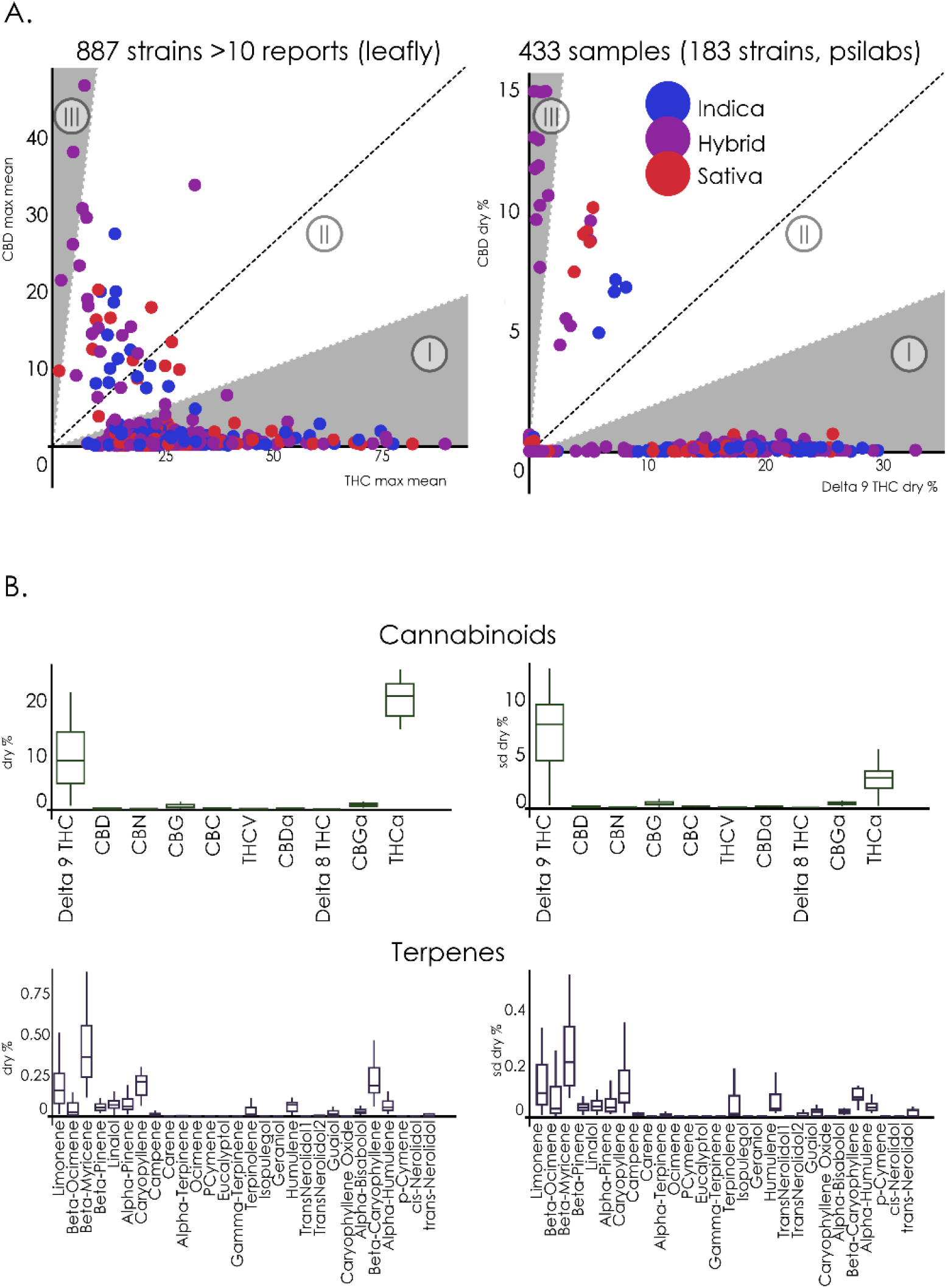
Chemical composition of cannabis strains. A. Scatter plot of CBD vs. THC (max mean content) for all strains (left panel) and for the 183 strains included in the PSI Labs dataset, by dry % (right panel). The background is divided by chemotype (THC:CBD ratios), where Chemotype I indicates 5:1, Chemotype III indicates 1:5, and Chemotype II is in the middle of both (Jikomes & Zoorob, 2018). While three different chemotypes could be identified, in both cases chemotype I (high THC content) was clearly overrepresented in the data. B. Cannabinoid and terpene content data extracted from PSI Labs. Left: boxplots of the mean dry content of 10 cannabinoids and 26 terpenes across multiple samples of the same strain. Right: the variability of the mean dry content across samples of the same strain (mean ± STD).

Fig. 7B shows the means and standard deviations of the compiled data for 10 cannabinoids and 26 terpenes across multiple samples of a strain included in the PSI Labs dataset. While some terpenes appeared to be robustly detected in the strain, the relatively large error bars indicated a considerable level of variability. Next, we addressed in more detail the association between cannabinoid content, terpene content, flavours, psychoactive effects, and cannabis species. For this purpose, each of the 183 strains in the PSI Labs dataset was described by a cannabinoid and terpene vector. We computed the Spearman correlation between these vectors to weight the links connecting the nodes that represented the strains. This resulted in cannabinoid and terpene similarity networks, which are shown in Fig. 8A and 8B, respectively. The network on the left panel of Fig. 8A is color-coded based on the application of the Louvain algorithm (Q = 0.041) to the cannabinoid similarity network, yielding a total of 8 modules, with the largest 3 represening ≈94% of the strains. This modular structure paralleled the classification into the three chemotypes.

**Fig. 8.**
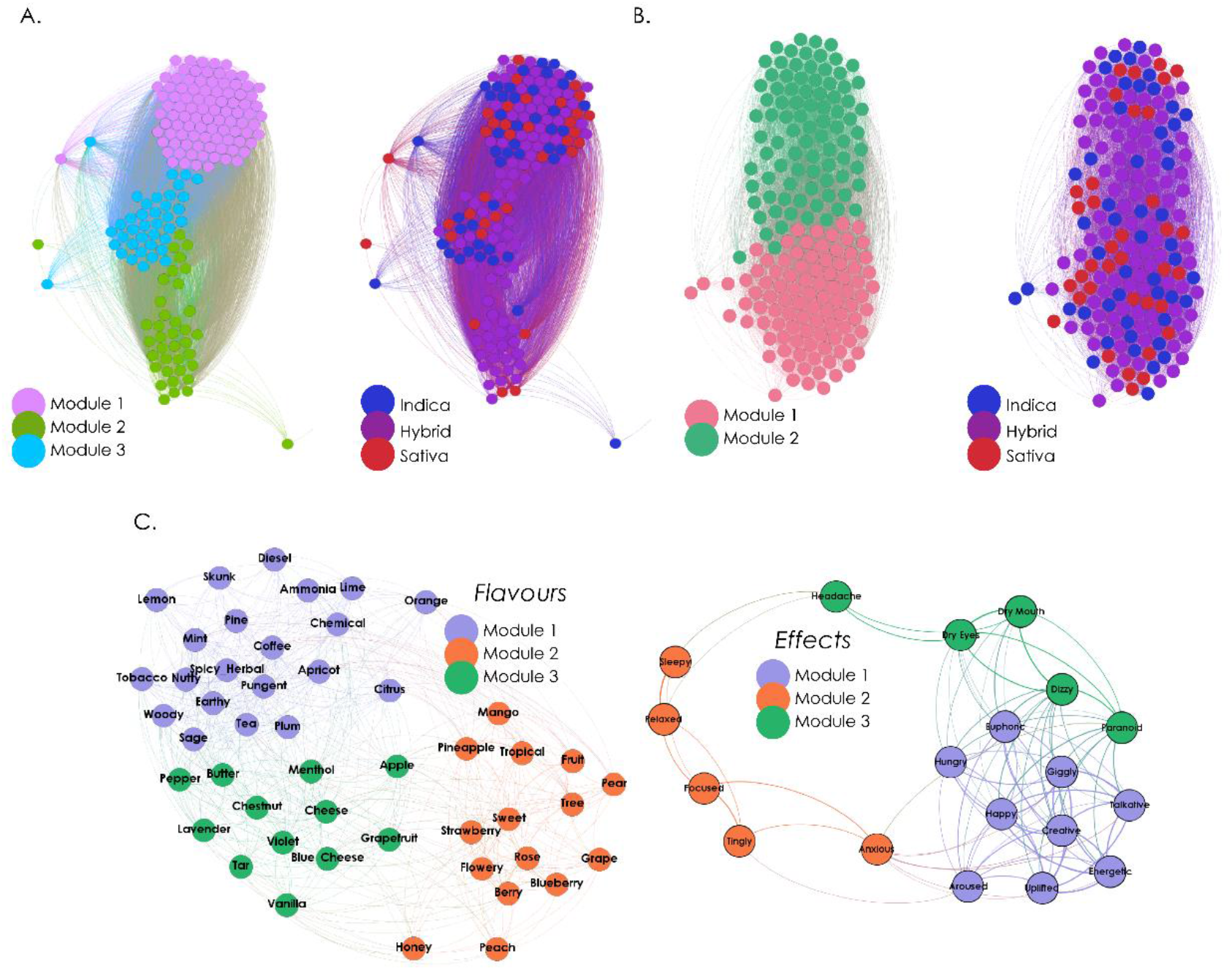
Association of strains, effect tags, and flavour tags in terms of chemical composition. A. Network of similarity in cannabinoid content. Each node represents a strain, and their spatial proximity is based on the correlation between their corresponding cannabinoid profiles. Nodes in the left panel are color-coded based on modularity analysis, while nodes in the right panel are color-coded based on species. B. Same analysis as in Panel A, but for the similarity in terpene content. C. The network in the left represents the association between flavour tags, based on the correlation of the terpene profiles averaged across all strains for which the corresponding flavour tags were reported. The network in the right presents the same analysis for effect tags.

The network on the right is color-coded based on cannabis species: the first and largest module contained strains belonging to all species (similar to chemotype I); another module, situated in the middle, presented a more balanced proportion of species, but also contained a smaller proportion of strains (similar to chemotype II), and the remaining module was composed mostly by hybrids (as in chemotype III). Since this classification used more information than the THC:CBD ratios, it corresponds to a multi-dimensional analogue of the standard chemotype characterization.

Fig. 8B shows the network obtained by correlating strains by their terpene vectors. The network on the left is color-coded based on the results of the Louvain algorithm (Q = 0.245), yielding only two modules. The network on the right is color-coded based on cannabis species. Since there are multiple terpenes in cannabis, without equivalents of main active cannabinoids such as THC and CBD, the chemical description in terms of terpenes is necessarily multi-dimensional. As with the semantic analysis of written reports, the association of strains based on the terpene profiles was bimodal and without a clear differentiation in terms of species.

Finally, we explored how effects and flavours were related based on the terpene content of the strains (Fig. 8C). We generated a terpene vector associated with each effect and flavour tag by averaging the terpene content across all the strains for which that tag was reported. The left panel of Fig. 8C shows how flavour tags (nodes) related in terms of the correlation of their associated terpene vectors (weighted links). Modularity analysis (Q = 0.324) yielded a module comprising intense and pungent flavours (“skunk”, “diesel”, “chemical”, “pungent”) combined with citric flavours (“lemon”, “orange”, “lime”, “citrus), a second module containing sweet and fruity flavours (“mango”, “strawberry”, “sweet”, “fruit”, “grape”), and a third module with a mixture of salty and sweet flavours (“cheese”, “butter”, “vanilla”, “pepper”). Modularity analysis (Q = 0.194) of the network of effect tags associated by terpene similarity (Fig. 8C, right panel) yielded three modules resembling the clustering of effects presented in Fig. 3C, where we found groups consisting of unwanted effects, stimulant effects and soothing effects, with an intermediate group associated with miscellaneous effects of smoked marihuana. Module 1 contained mostly stimulant effects (“energetic”, “euphoric”, “creative”, “talkative”, among others), module 2 contained soothing effects (“sleepy”, “relaxed”), and module 3 contained unwanted effects such as “headache”, “dizzy”, “paranoid” (with the exception of “anxious”, which was included in module 2). The fact that the network of effects associated by terpene content similarity reflected the hierarchical clustering of effects obtained from flavour association (Fig. 3C) reinforces the link between flavours and the psychoactive effects of cannabis.

## Discussion

We presented a novel synthesis of multi-dimensional data on a large number of cannabis strains with the purpose of developing an integrated view of the relationship between psychoactive effects, perceptual profiles (flavours) and chemical composition (terpene and cannabinoid content). As a result of this analysis, we established that cannabis species can be inferred from self-reported effect and flavour tags, as well as from unstructured written reports, which also revealed that topics associated with subjective effects and therapeutic use had different prevalence in indicas compared to sativas. This classification was obtained using supervised (random forests) and unsupervised (network modularity maximization) methods. As suggested by the previous literature (Casano et al., 2011; Fischedick J, 2015; Pollastro, Minassi, & Fresu, 2018), we found that classifiers based on the reported flavours achieved surprisingly high accuracy in the classification of strains. Furthermore, flavour and effect tags did not manifest independently, but presented significative correlations that were expected from studies showing synergetic effects between cannabinoids and terpenes (Russo, 2011, 2019). Finally, in spite of high variability in the reported chemical compositions, we corroborated the presence of expected flavour-terpene associations. In the following, we discuss the implications of our work in the context of leveraging large volumes of data produced under naturalistic conditions in combination with quantitative chemical analyses for the classification and characterization of cannabis strains.

The practical relevance of our results is manifest in the need to develop fast, cheap and reliable methods for cannabis strain and species identification. Over the past years, marihuana species (indica / sativa) have been challenged by the scientific community as reliable markers of the effects elicited by the consumption of the plant (Piomelli & Russo, 2016; Pollastro et al., 2018; Russo, 2019), pointing towards objective chemotype descriptors (mainly THC:CBD ratios) as a new golden standard. According to this characterization, THC is often considered the active compound related to many of the negative outcomes of cannabis consumption (Volkow et al., 2016), while CBD (or combinations of CBD and THC) is associated with most of the purported medicinal properties (Fogaça, Campos, Coelho, Duman, & Guimarães, 2018; Hahn, 2018; Hurd et al., 2019; Lorenzetti, Solowij, & Yücel, 2016; Nadulski et al., 2005; Nuutinen, 2018; Russo, 2019; Vandrey et al., 2015). Although there is no doubt that a precise chemical description of the plant is the most accurate and reliable predictor of the elicited effects, this approach is likely impractical in the present market (Nie et al., 2019). In the first place, this approach requires technology for quantitative chemical analysis that is beyond the reach of many dispensaries and growers. Furthermore, variations in the concentration of cannabinoids are high even within the same strain, depending on factors such as age, environmental conditions, generation, and geographical location (Casano et al., 2011; Fetterman et al., 1971; Nuutinen, 2018; Russo, 2011). Finally, the predictive value of chemotypes has been questioned in markets where consumers increasingly demand higher THC content (Freeman et al., 2019, 2018; Jikomes & Zoorob, 2018; Smart, Caulkins, Kilmer, Davenport, & Midgette, 2017).

Our results suggest that perceptual profiles (reported flavours) and terpene quantification show merit for the characterization of cannabis strains. Both tagged psychoactive effects and perceived flavours were capable of predicting species with high accuracy. Terpenes are highly conserved across generations (Aizpurua-Olaizola et al., 2016; Casano et al., 2011), can be synergetic with cannabinoids (Russo, 2011), and have psychoactive properties by themselves (Nuutinen, 2018). It follows from our analysis that users could count on perceptual faculties to select strains when seeking specific effects. Further research in controlled laboratory settings is required to test the capacity for predicting psychoactive effects based on sensory information.

There is increasing evidence that the subjective and therapeutic effects of cannabis are a result of the synergy between a diverse group of active ingredients which include THC and CBD, alongside other cannabinoids and terpenes (Baron, 2018; Nuutinen, 2018; Russo, 2011). This observation supports the need for a multi-dimensional characterization that does not neglect terpene content, and therefore the associated flavours. We found that, even with overall high levels of THC across all strains (Jikomes & Zoorob, 2018), the subjective experiences reported by users were capable of clustering strains by species, not only based on effects but also on the reported flavours. Moreover, the clustering of strains with names evoking certain flavours, even while not botanically validated (Clarke & Merlin, 2013), supported that terpene content is a well-preserved property in the strains.

As in the sativa-indica gradient revealed by the analysis of effect and flavour tags, the semantic analysis of unstructured written reports clearly captured the distinction between “sativa like” and “indica like” strains. It is interesting to note that, in spite of overall high THC content across all strains, the specific words that emerged from LSA topic detection applied to reports of sativas and indicas represented a large proportion of positive and desired effects, such as relaxing and uplifting effects (Corral, 2001). Natural language analysis also established that, even while therapeutic and subjective effects were thoroughly discussed throughout the written reports, seed acquisition and plant growing were also prominently featured.

Concerning terpene and cannabinoid profiles, it is important to note that cannabinoids showed a clear trimodal structure, in accordance with the three chemotypes described by Jikomes & Zoorob (Jikomes & Zoorob, 2018), which were obtained based only on THC and CBD concentrations. The fact that a trimodal grouping of the strains was also obtained based on 10 cannabinoids could imply that the complex interactions of a larger number of active molecules might project into a reduced tri-dimensional space without significant loss of information. The concept of multi-dimensional chemotype should be further explored in controlled laboratory conditions to develop more accurate objective descriptors of different cannabis strains and their elicited effects. Conversely, strains were organized bimodally by their terpene content. This observation is interesting in the context of the flavour-effect associations identified in our work, which were essentially organized into two groups: stimulating and sedative effects. These associations add support to the active role of terpenes (Nuutinen, 2018). The analysis of effect association via terpene content similarity yielded results convergent with those obtained from correlating flavour and effect tags, adding further support to psychoactive effects being mediated by terpenes and/or their synergy with cannabinoids.

The strengths of our study stem from the analysis of large volumes of data impossible to obtain under controlled laboratory conditions, but this approach also leads to limitations inherent to self-reporting by users, preventing objective verification of the consumed strains, as well as their doses. Limitations are also inherent to the assumption of cannabis strains as having homogeneous chemical compositions. Previous work showed that cannabinoid content can present ample variation within single strains (Fischedick J, 2015; Jikomes & Zoorob, 2018), and our results show that similar considerations apply to terpene profiles. Future studies could address a smaller sample of strains more thoroughly investigated in terms of their chemical composition, thus allowing the study of correlations between self-reported psychoactive effects, flavours, and environmental factors that could impact on cannabinoid and terpene content.

## Conclusions

After decades of prohibition, the legal cannabis industry for therapeutic and recreational use is growing at a quick pace, but nevertheless it is at its infancy. Considerable evidence suggests that commercially available strains are highly variable in their chemical composition and psychoactive effects. In comparison, more mature industries, such as that of winery, have arrived to reliable standards (e.g. Merlot, Cabernet, Syrah) that are trusted and understood by the consumers. By extracting information from different sources of data, our work suggests that the development of standards in the cannabis industry should not only focus on psychoactive effects and cannabinoid content, but also take into account scents and flavours, which constitute the perceptual counterpart of terpene and terpenoid profiles.

## Additional file

Additional file 1: supplementary tables with details on cannabis strains, flavour and effect tags, and supplementary analyses to support the results.

## Abbreviations

THC: Tetrahydrocannabinol
CBD: Cannabidiol
AUC: Area under the receiver operating characteristic curve
LSA: Latent Semantic Analysis
SVD: Singular Value Decomposition
PCA: Principal Component Analysis
FDR: False Discovery Rate.

## Acknowledgments

We thank the founders, curators and contributors of Leafly and PSI Labs for publicly sharing the datasets used in this study.

## Authors’ contributions

CP and ET conceived the study. AF and CP performed the analyses and elaborated all figures. ASF, FC and FZ provided technical assistance, contributed towards interpreting the results, and gave feedback on early versions of the manuscript. AF, CP and ET wrote the final version of the manuscript.

## Availability of data and materials

The datasets supporting the conclusions of this article are available in the Mendeley Data repository (DOI: 10.17632/6zwcgrttkp.1, https://tinyurl.com/yyzkk78r).

## Ethics approval and consent to participate

The authors did not perform experiments nor acquired data other than information already available in publicly shared websites. This study only provides results from statistical analyses and does not contain information that could lead towards the identification of users. Details concerning confidentiality, terms of use and copyright can be found in the Leafly website: https://www.leafly.com/company/privacy-policy

## Consent for publication

Not applicable.

## Footnotes

1 We follow the notation where 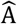refers to a matrix and A_ij_ to a particular entry.

2 As a naming convention, we identified sets of strains with a flavour in their name using that flavour, e.g. the group of “lemons” comprised strains such as Lemon Skunk and Lemon Diesel. We also described these groups by their general category, e.g. “lemons”, “grapefruits”, “strawberries” were labeled “fruits”.

